# BacPROTACs outperform inhibitors in *Mycobacterium tuberculosis*

**DOI:** 10.64898/2026.06.12.731830

**Authors:** David A. Brown, Joshua J. Davies, Selina Fecht, Yujia Zhang, Simone Kunzelmann, Lois Kent, Mark Skehel, Francesca E. Morreale

## Abstract

Antibiotic discovery has long relied on occupancy-driven inhibition, leaving a vast number of potential bacterial targets undrugged.^1^ Targeted protein degradation offers a mechanistically distinct alternative to inhibition, yet its application to antibacterial drug discovery remains largely unexplored.^2–4^ Here we describe the development of first-in-class heterobifunctional bacterial proteolysis targeting chimeras (BacPROTACs) directed against an essential *Mycobacterium tuberculosis* protein, 4’-phosphopantetheinyl transferase (PptT).^5^ Leveraging the modular architecture of BacPROTACs, we repurposed PptT inhibitors by incorporating them into degraders, yielding compounds with markedly improved antimycobacterial activity. Integrating *in vitro* and cellular approaches, we developed a characterisation pipeline to assess protein degradation in bacteria, applicable to future BacPROTAC programmes. Our study establishes targeted protein degradation as a strategy for antibacterial drug discovery.

## Introduction

Targeted protein degradation (TPD) is an emerging drug discovery paradigm whereby a protein of interest is recruited to the cellular proteolytic machinery for selective elimination.^2, 6–8^ In oncology, substantial academic and industrial investment has advanced the field at remarkable pace, culminating in the recent landmark FDA approval of the first PROTAC degrader.^9^ However, the translation of this strategy to antibacterial drugs remains underexplored, due to the fundamental differences in protein degradation pathways between eukaryotic and prokaryotic cells.^10, 11^ Progress has been further hampered by the limited number of validated ligands of bacterial protein degradation effectors, reflecting the historically poor investment in antibacterial drug discovery relative to human protein targets.^12^ Nevertheless, TPD offers a transformative strategy to expand the druggable bacterial proteome, diversify antibiotic mechanisms of action, and circumvent existing resistance mechanisms. To this end, early progress has focused on proof-of-concept BacPROTACs that concurrently bind target proteins and the mycobacterial ClpC1P1P2 proteolytic complex.^4^ Clp proteases are the primary effectors of protein quality control in bacteria,^13, 14^ functioning as the prokaryotic counterpart to the eukaryotic ubiquitin-proteasome system, and are thus amenable to reprogramming by degrader molecules. ClpC1P1P2-redirecting BacPROTACs demonstrated the feasibility of proximity-induced protein degradation in bacteria, yet initial validation relied on heterologous model proteins, and on ClpC1 autodegradation.^4, 15–17^

Building on these foundational studies, we aimed to develop heterobifunctional BacPROTACs by repurposing ligands of essential mycobacterial proteins. Although numerous ligands targeting bacterial proteins have been identified, most failed at different stages of development due to insufficient cellular activity, limited membrane permeability, or inadequate selectivity between host and pathogen proteins.^18, 19^ The modular structure of BacPROTACs enables the repurposing of such ligands into degrader warheads, fundamentally altering their permeability and mechanism of action, with the potential to confer host-pathogen selectivity and retain activity against inhibitor-resistant mutants. This approach unlocks a novel therapeutic modality, without the need for novel chemical matter.

To explore this potential, we focused on *Mycobacterium tuberculosis* (*Mtb*), which remains one of the leading causes of death from a single infectious agent worldwide.^20^ The expanding prevalence of multidrug-resistant and extensively drug-resistant strains further complicates the standard six-month treatment regimen, necessitating more complex multidrug combinations and underscoring the urgent need for antitubercular agents with novel mechanisms of action.^21, 22^ We harnessed inhibitors of 4’-phosphopantetheinyl transferase (PptT),^23–25^ an essential enzyme for the survival of *Mtb* both *in vitro* and in mice,^5, 26^ to develop BacPROTACs that recruit PptT to the ClpC1P1P2 proteolytic complex for targeted degradation (Fig. 1a). We report the first two series of BacPROTACs derived from repurposed mycobacterial inhibitors with improved antitubercular activity.

**Figure 1.**
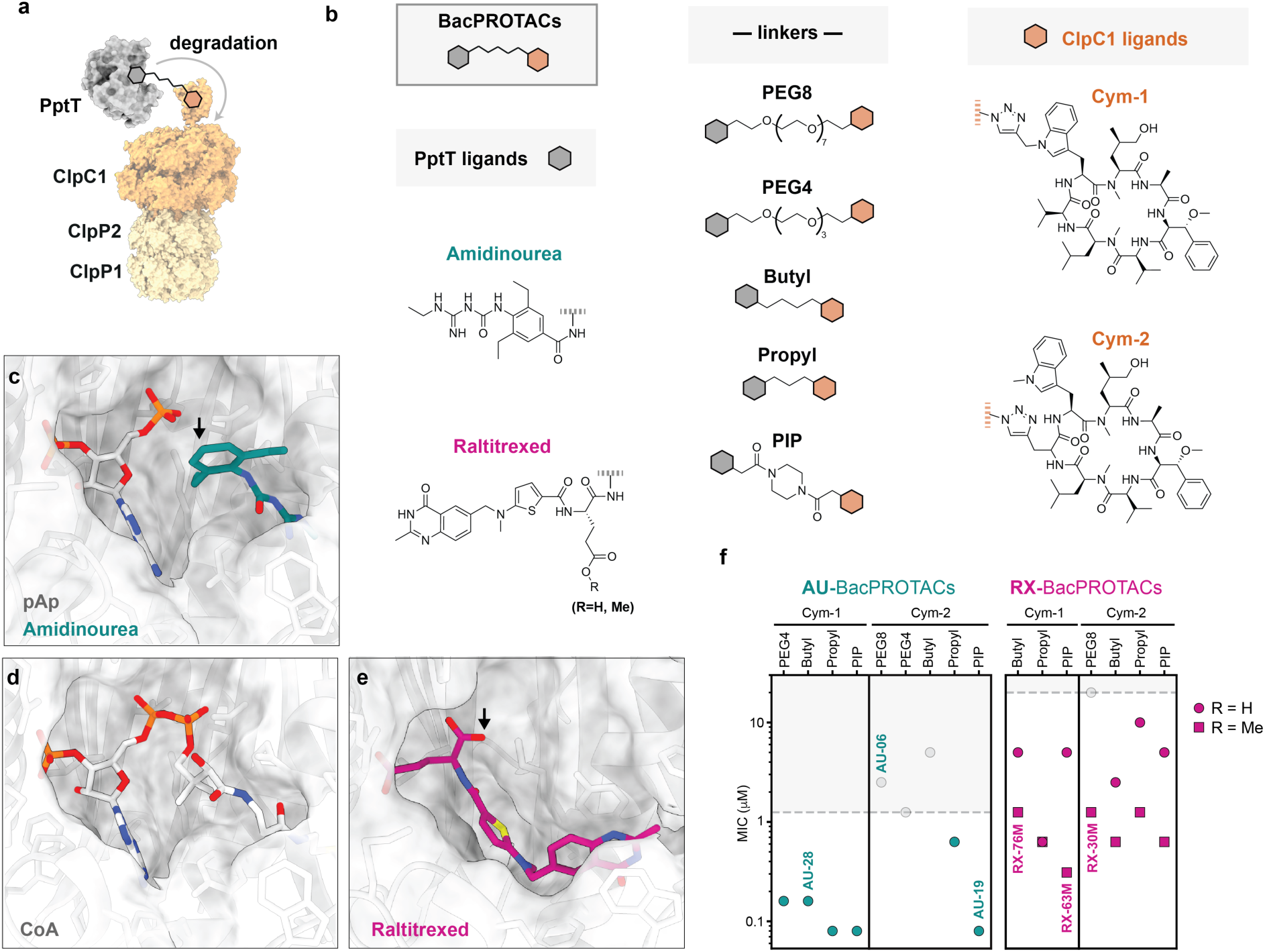
BacPROTAC design and antimycobacterial activities. **(a)** Schematic of the targeted protein degradation approach. The BacPROTAC degrader recruits the essential mycobacterial protein 4’-phosphopantetheinyl transferase (PptT) to the ClpC1P1P2 proteolytic complex. **(b)** Heterobifunctional BacPROTACs synthesised in this study, composed of Amidinourea (AU)- or Raltitrexed (RX)-based ligands engaging PptT,^23, 24^ two cyclomarin ligands engaging ClpC1 (Cym-1, Cym-2),^16^ and five different linkers. Dotted lines indicate linker connection points; grey and orange hexagons represent PptT and ClpC1 ligands, respectively. Crystal structures of PptT bound to: **(c)** AU 8918^23^ + pAp (PDB: 7N8M),^25^ **(d)** CoA (PDB: 4U89),^35^ and **(e)** Raltitrexed (PDB: 8GKF) ^24^ visualised using UCSF ChimeraX.^36^ Protein-ligand structures enable designing the optimal linker attachment point (black arrow) and show that AU and RX ligands occupy distinct portions of the CoA binding site. **(f)** Plotted MIC screening results of synthesised BacPROTACs against *Mycobacterium tuberculosis* H37Rv (Extended Data Fig. 1d). MIC is defined as the lowest concentration inhibiting bacterial growth by 90% relative to DMSO (100% growth) and rifampicin (0% growth) controls. Dashed lines indicate the MIC of the free ligand combinations for each BacPROTAC series. Compounds with MIC values exceeding the corresponding threshold are shown in grey in the shaded area. BacPROTACs selected for further characterisation are indicated by their compound names. For the RX series, methylated compounds (R = Me) serve as prodrugs to improve membrane permeability. The MIC screening shows one representative experiment of *n* = 2 biological replicates performed in technical quadruplicate.

To date, the fundamental differences between eukaryotic and prokaryotic protein degradation pathways have left the field without established guidelines for the mechanistic characterisation of degraders in bacteria. Here, we developed an integrated platform combining *in vitro* and cellular assays tailored to mycobacteria, to distinguish *bona fide* degraders from non-degraders. We anticipate that this work will serve as a blueprint for future BacPROTAC development and pave the way for TPD in antibiotic discovery.

### Incorporating PptT inhibitors into degraders yields effective antitubercular compounds

To investigate whether repurposing inhibitors of *Mtb* proteins as degrader warheads could improve whole-cell activity, we designed two series of PptT-targeting BacPROTACs (Fig. 1b). PptT catalyses the covalent transfer of the 4’-phosphopantetheine moiety from coenzyme A (CoA) to acyl carrier protein domains.^5, 27^ This post-translational modification is a prerequisite for the synthesis of complex lipids that constitute the mycobacterial cell envelope, which underpins both *Mtb* virulence and restricted drug permeability.^5, 28, 29^ Recent studies have identified two classes of small-molecule PptT inhibitors with distinct chemical scaffolds targeting different regions of the active site (Fig. 1c-e). Amidinourea (AU)-based compounds bind the pocket occupied by the phosphopantetheine arm of CoA, inhibit catalysis, and show whole-cell activity against *Mtb*.^23, 25^ In contrast, Raltitrexed (RX), an inhibitor of human thymidylate synthase recently found to target PptT *in vitro*, fully displaces CoA and inhibits catalysis but lacks whole-cell activity, reportedly due to efflux.^24^ By incorporating both RX- and AU-based inhibitors into degraders, our study aimed to improve cellular activities while comprehensively evaluating how degrader architecture, including ligand choice, linker length and attachment points, influence ClpC1P1P2-mediated degradation in *Mtb*. To this end, we synthesised BacPROTACs composed of five different linkers and two previously reported Cyclomarin derivatives for ClpC1 engagement,^4, 16^ bearing distinct exit vectors for linker attachment (Cym-1 and Cym-2, Fig. 1b). Cym-1 and Cym-2 have been used in previous proof-of-concept studies to engage the N-terminal domain of ClpC1 (ClpC1_NTD_) and recruit target proteins for unfolding by the hexameric AAA+ ATPase rings of ClpC1, which subsequently translocate the unfolded polypeptide into the ClpP1P2 proteolytic chamber for degradation.^4,13, 30, 31^

We synthesised a total of 23 different BacPROTACs including 9 AU-based, 7 RX-based degraders with free carboxylic acid and 7 corresponding methyl esters (Fig 1b, Extended Data Fig. 1a-c), previously found to improve cellular uptake, and to be metabolised back to the free carboxylic acid once inside the cell.^24, 32^

To assess whether the synthesised bifunctional molecules displayed improved whole-cell activity, we screened the 23 BacPROTACs for minimum inhibitory concentrations (MIC) against *Mtb* H37Rv and compared BacPROTACs to the corresponding free ligand combinations (Fig. 1f, Extended Data Fig. 1d). We also confirmed that representative compounds were well tolerated in human cell lines (Extended Data Fig. 1e). Comparison of BacPROTAC and free-ligand MICs enriches for compounds for which protein degradation is advantageous over inhibition, although MIC differences among BacPROTACs may reflect a combination of factors including cell permeability, necessitating further mechanistic validation.

MIC screening revealed that, similarly to eukaryotic PROTACs,^33, 34^ varying ligands, linkers and linker attachment sites critically shapes the structure-activity relationship (SAR), resulting in large differences in cellular potencies. We found that the AU series was generally more potent than the RX series. Crucially, most BacPROTACs outperformed the corresponding free ligand combinations, with compounds from the AU series showing MICs as low as 80 nM (Fig. 1f, Extended Data Fig. 1d).

Collectively, the MIC results highlighted the need to systematically probe linker-ligand combinations to optimise BacPROTAC potency beyond that of the parent inhibitors and prompted further investigation into the mechanistic basis of BacPROTAC activity.

### PptT-directed BacPROTACs function as proximity-inducing degraders *in vitro*

To determine whether BacPROTACs were functioning through a degrader mechanism of action, and to rationalise the efficacy trends observed in the MIC screening, we selected a representative set of compounds for *in vitro* characterisation (Fig. 2a, Extended Data Fig. 2a). In order to capture both the rate and extent of PptT degradation induced by different BacPROTACs, we set up an assay monitoring degradation of fluorescently labelled recombinant *Mtb* PptT (PptT^ATTO488^) by the reconstituted *Mtb* ClpC1P1P2 complex. In parallel, we assessed BacPROTAC-mediated ternary complex formation using PptT^ATTO488^ and ClpC1_NTD_ in a fluorescence polarisation (FP) assay (Fig. 2b).

**Figure 2.**
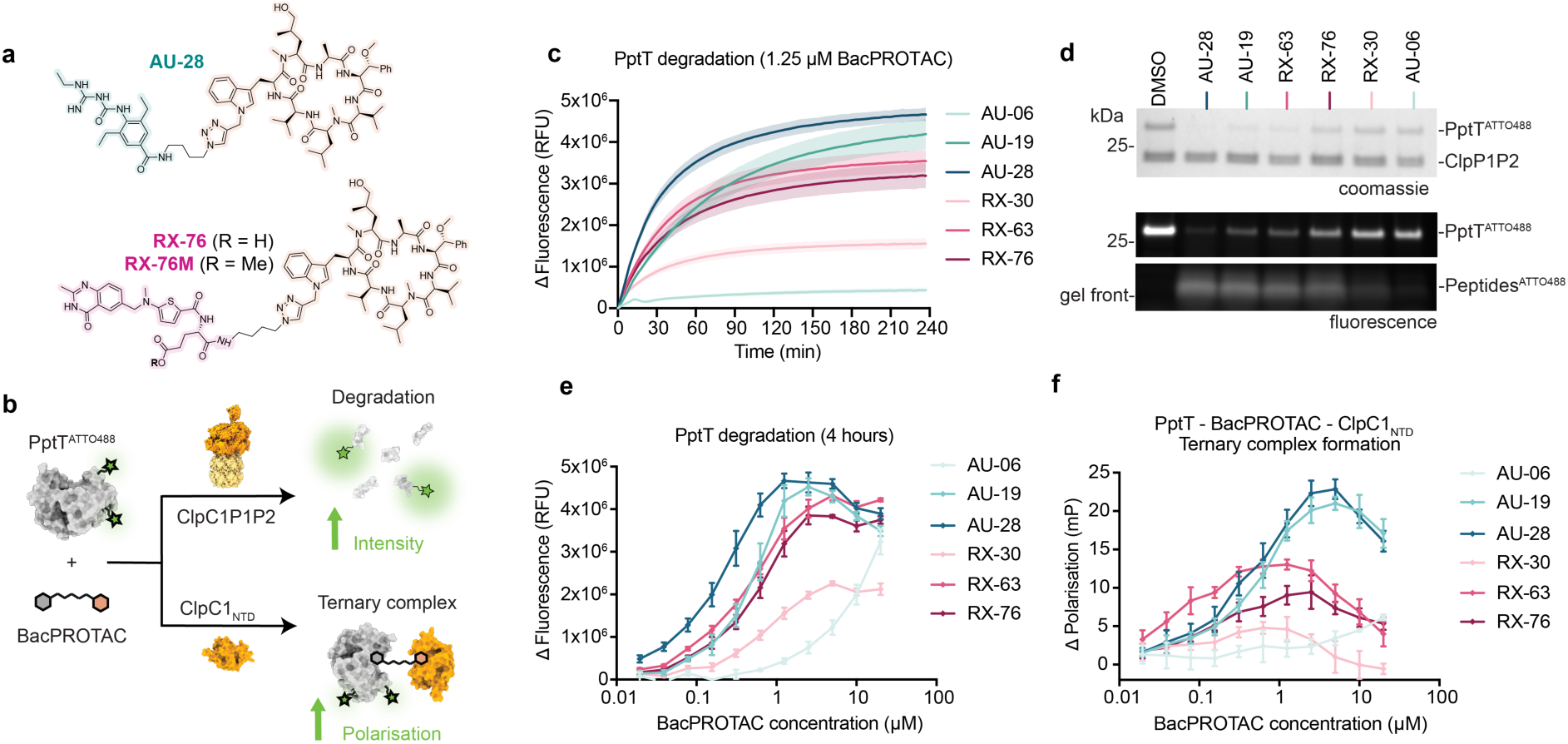
BacPROTACs mediate PptT degradation through ternary complex formation. **(a)** Chemical structures of the representative BacPROTACs AU-28, RX-76 and the corresponding prodrug RX-76M. **(b)** Schematic of the *in vitro* strategy to characterise BacPROTAC-mediated PptT degradation and ternary complex formation. Fluorescently labelled PptT^ATTO488^ is used to measure degradation by increased fluorescence intensity upon ClpC1P1P2-mediated proteolysis of the folded protein into peptides (dequenching), and ternary complex formation by increased fluorescence polarisation (FP) upon binding of BacPROTAC and ClpC1_NTD_ to PptT^ATTO488^. **(c)** Kinetics of PptT^ATTO488^ degradation by six BacPROTACs at 1.25 µM, revealing differences in maximal degradation (D_max_) and initial rate depending on linker length and type. Fluorescence intensity values are normalised by subtraction of t = 0 and DMSO background. Data are mean ± s.d. (shaded area). **(d)** SDS-PAGE analysis of the fluorescence degradation assay endpoint (t = 4 h) shown in **(c)**, imaged by in-gel fluorescence and Coomassie staining. Folded PptT is degraded into fluorescent peptides running at the gel front, corroborating the trends observed by fluorescence intensity readout. **(e)** Concentration-dependent PptT degradation by BacPROTACs, quantified as the difference in fluorescence intensity between t = 0 and the assay endpoint (t = 4 h) at each concentration tested, after DMSO background subtraction. **(f)** FP dose-response of BacPROTACs titrated into PptT^ATTO488^ and ClpC1_NTD_, indicating ternary complex formation with a hook effect at higher concentrations. Data in **(c)**, **(e)**, **(f)** are mean ± s.d. (*n* = 4 technical replicates; one representative experiment of *n* = 3 independent experiments shown).

In these assays, BacPROTACs induced ternary complex formation and PptT^ATTO488^ degradation with markedly different efficiencies (Fig. 2c-f, Extended Data Fig. 2b,c). AU-28 showed the fastest initial degradation rate (V_0_) and highest maximal degradation (D_max_), whereas long-linker BacPROTACs displayed considerably lower PptT degradation, consistent with negligible ternary complex formation. Notably, the conformationally constrained linker of AU-19 yielded comparable ternary complex formation to AU-28 but slower degradation, suggesting that linker rigidity and attachment site influence the orientation of PptT relative to the ClpC1 unfoldase pore, constraining productive degradation. PptT^ATTO488^ degradation and ternary complex formation displayed the characteristic ‘hook effect’,^37^ whereby binary saturation of PptT and ClpC1 at super-stoichiometric compound concentrations reduced ternary complex formation and thus degradation (Fig. 2e,f). Finally, addition of an excess of the free ligands (Cym-1, AU-08, RX) reduced degradation, confirming that BacPROTAC-mediated degradation depends on bivalent engagement of PptT and ClpC1 (Extended Data Fig. 2d-g). Of note, we found that linker length has a larger impact on the effective concentration range in AU-BacPROTACs than in RX-BacPROTACs. Combined with the reduced ability of free ligands to outcompete AU-BacPROTAC-mediated degradation, this suggests positive cooperativity in the AU-BacPROTAC binding mode. (Extended Data Fig. 2d,e,h).

Taken together, these results demonstrate that BacPROTACs function as proximity-inducing degraders of PptT *in vitro*. Despite showing interesting SAR trends, *in vitro* degradation profiles did not fully correlate with measured MICs. In particular, the considerably different D_max_ achieved by RX-76 and RX-30 (Fig. 2c–e) contrasted with their equal MICs (Fig. 1f), prompting investigation into the cellular basis for this discrepancy.

### Proteomic analysis uncovers distinct growth inhibition mechanisms

Given the limited correlation between *in vitro* degradation activity and MIC amongst RX-BacPROTACs, we sought to investigate the mechanistic basis for the observed antitubercular activity.

To determine whether antimycobacterial activity correlates with PptT degradation in cells, we analysed the proteomic changes associated with the *Mtb* growth inhibition phenotype (Fig. 3a). We selected one representative BacPROTAC from each series (AU-28 and RX-76M, composed of the same linker and Cym ligand) and performed proteomic analysis utilising concentrations along the growth inhibition curve, spanning intermediate to complete inhibition phenotypes (Fig. 3b). Proteomics analysis revealed that AU-28 induced dose-dependent PptT degradation at concentrations as low as 40 nM, reaching the highest fold change at MIC (Fig. 3c,d). Degradation was specific to PptT and increased over time (Extended Data Fig. 3a). As a control, we performed a similar analysis for the combination of free ligands composing AU-28 and confirmed that no degradation is observed when the two ligands are not connected via a linker (Extended Data Fig. 3b,c). In sharp contrast to AU-28, RX-76M did not induce detectable PptT degradation at any of the concentrations tested (Fig. 3e), suggesting that the antitubercular activity of RX-BacPROTACs is not driven by PptT degradation, and accounting for the lack of correlation between *in vitro* degradation efficiency and MIC.

**Figure 3.**
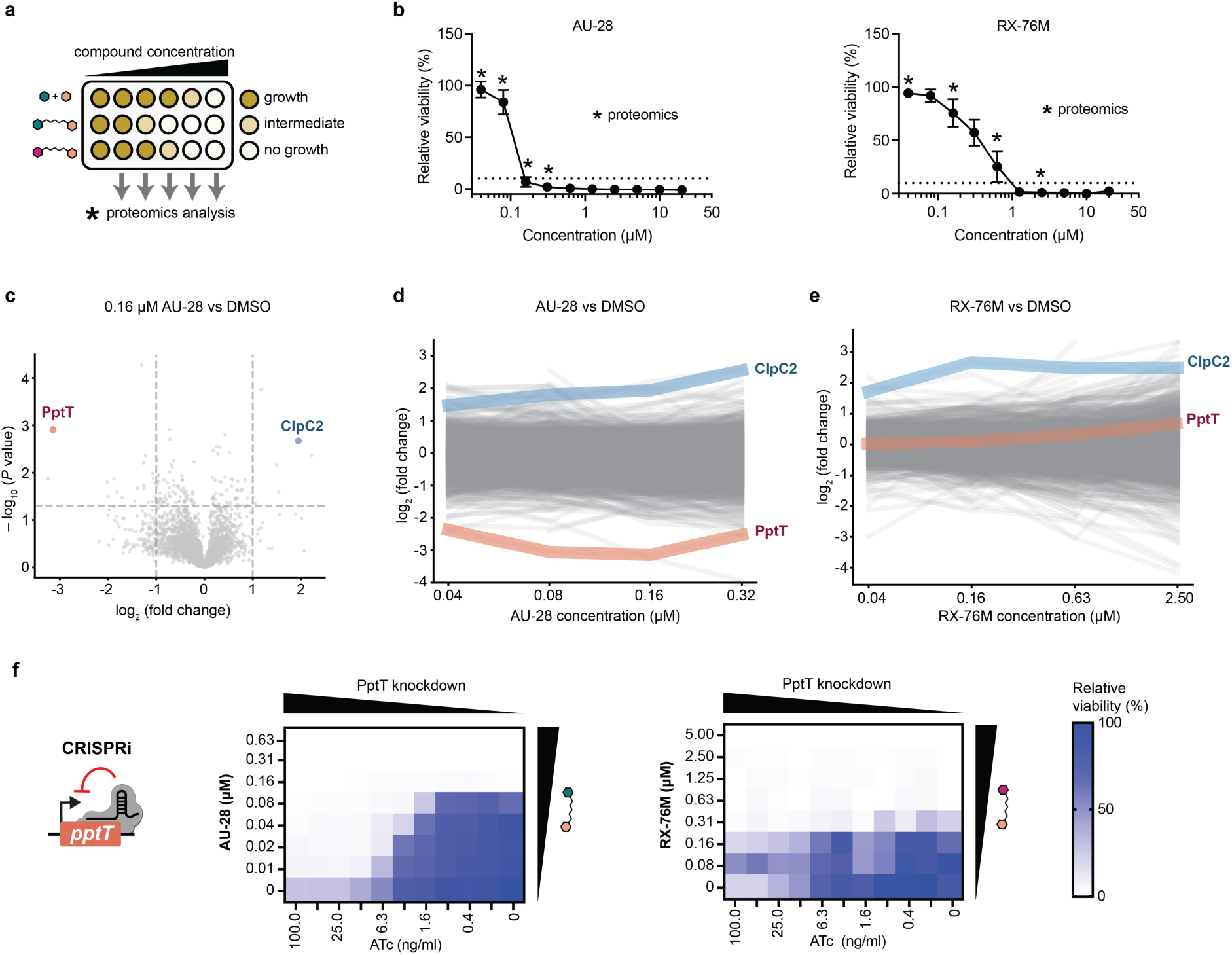
Cellular characterisation of BacPROTAC mechanisms. **(a)** Schematic of the proteomic strategy, in which *Mycobacterium tuberculosis* cultures were treated with a range of BacPROTAC concentrations. Conditions inducing intermediate to complete growth inhibition were selected to analyse proteomic changes associated with the growth inhibition phenotype. **(b)** *Mtb* growth inhibition induced by AU-28 (left) or RX-76M (right) treatment. The dotted line indicates the minimum inhibitory concentration (MIC) threshold (90% growth inhibition), relative to DMSO (100% growth) and rifampicin (0% growth) controls. Asterisks indicate BacPROTAC concentrations selected for proteomic analysis. Data are mean ± s.d. of *n* = 4 technical replicates; one representative of *n* = 4 independent experiments. **(c)** Volcano plot of proteomic changes in *Mtb* treated with 0.16 µM AU-28 relative to DMSO. Dotted lines indicate significance thresholds (– log_10_ (*P* value) = 1.3, log₂(fold change) = ± 1). PptT is significantly degraded and ClpC2 upregulated, consistent with previous observations upon cyclomarin treatment.^15, 39^ **(d)** Proteomic changes across AU-28 and **(e)** RX-76M concentrations. AU-28 induces selective PptT degradation and ClpC2 upregulation (*n* = 3 biological replicates, each performed in technical triplicate), while RX-76M leads to ClpC2 upregulation and no detectable PptT degradation (*n* = 2 biological replicates, each performed in technical triplicate). **(f)** CRISPRi-mediated transcriptional repression of *pptT* in *Mtb* modulated by increasing concentrations of anhydrotetracycline (ATc). The heat map shows a checkerboard assay in which ATc and BacPROTAC (left: AU-28, or right: RX-76M) are co-titrated. *Mtb* growth inhibition is normalised to vehicle (100% growth) and rifampicin (0% growth) controls. Depletion of PptT sensitised bacteria to AU-28 in an ATc dose-dependent manner, while RX-76M potency was unaffected. Data are mean of *n* = 3 technical replicates; one representative of *n* = 2 independent experiments shown.

To further dissect the mechanistic basis of antitubercular activity, we employed CRISPRi-mediated transcriptional repression of *pptT* in *Mtb*.^38^ We modulated knockdown by titrating increasing concentrations of the inducer anhydrotetracycline (ATc) and assessed sensitivity to AU-28 and RX-76M. Partial depletion of PptT hypersensitised bacteria to AU-28 in an ATc dose-dependent manner (Fig. 3f, Extended Data Fig. 3d), confirming on-target activity. In contrast, RX-76M potency was unaffected by PptT knockdown, indicating that the compound inhibits *Mtb* growth independently of PptT, neither through degradation nor inhibition, and thus acts through a distinct mechanism.

These results establish AU-28 as the first heterobifunctional degrader of an essential mycobacterial protein. Yet the absence of cellular PptT degradation by RX-76M contrasted sharply with our *in vitro* findings, prompting us to investigate the basis for this discrepancy. The lack of Raltitrexed whole-cell activity was previously attributed to efflux.^24^ However, our proteomics experiments suggest that the corresponding BacPROTAC RX-76M, similarly to AU-28, reaches a sufficient intracellular concentration to trigger ClpC2 upregulation (Fig. 3e), a signature response to Cym-based compounds.^15, 39, 40^ This observation makes efflux alone an unlikely explanation for the differences in cellular PptT degradation between the two compounds.

### Ligand binding mode underlies target engagement and degradation profiles

To rationalise the lack of cellular degradation by RX-76M, we hypothesised that CoA, which binds with high affinity to PptT^28^ and is a ubiquitous intracellular cofactor^41^, may compete with BacPROTACs for target engagement inside mycobacteria. This competition would differentially affect the two degrader series: while Amidinourea compounds must only displace the phosphopantetheine arm of CoA and are compatible with concurrent binding of the reaction product pAp,^23^ Raltitrexed must displace CoA across the entire active site at all stages of the catalytic cycle (Fig. 1c-e). ^24^ To test this hypothesis, we titrated CoA or pAp into a fixed concentration of AU-28 or RX-76 in an *in vitro* degradation assay (Extended Data Fig. 4a). Notably, we found that in the presence of CoA, degradation mediated by RX-76 became negligible, whereas AU-28 retained appreciable degradation activity (Fig. 4a). pAp even increased the initial rate of AU-28-mediated degradation, likely by displacing co-purified CoA, thereby priming PptT for efficient BacPROTAC engagement (Extended Data Fig. 4b,c). These findings were corroborated by FP measurements of ternary complex formation (Fig. 4b) and are collectively consistent with the distinct binding modes of the two ligands.^23, 24^

**Figure 4.**
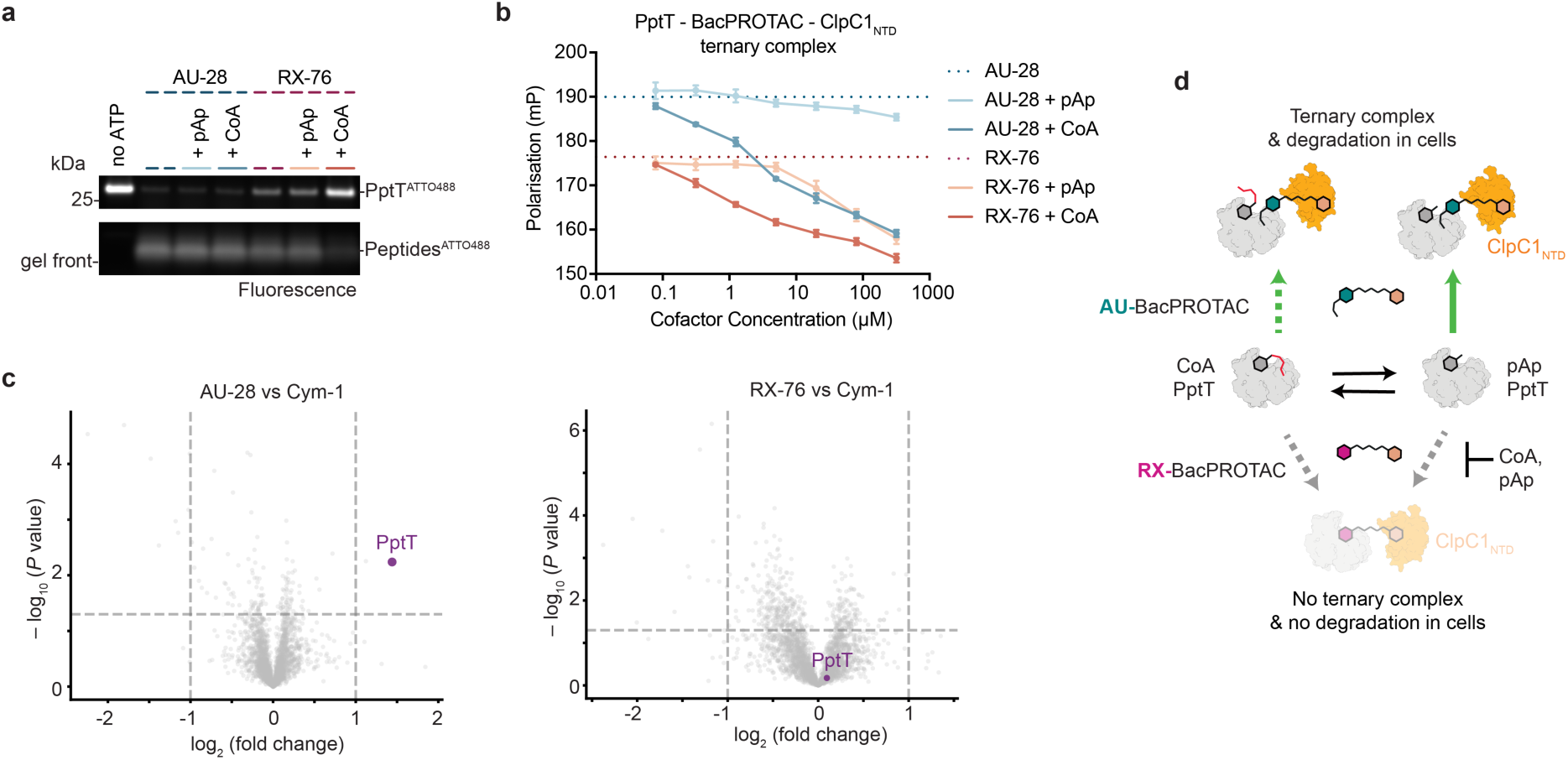
Cellular cofactors differentially modulate BacPROTAC engagement. **(a)** PptT ^ATTO488^ degradation induced by 1.25 µM AU-28 or RX-76 in the presence of 80 µM CoA or pAp, assessed at the assay endpoint (t = 4 h) by SDS-PAGE in-gel fluorescence (kinetic read in Extended Data Fig. 4b) **(b)** Fluorescence polarisation dose-response of CoA or pAp titrated into 1.25 µM BacPROTAC, PptT^ATTO488^ and ClpC1_NTD_, revealing differential BacPROTAC susceptibility to cofactor-induced ternary complex dissociation. Data in **(a, b)** are mean ± s.d. (*n* = 4 technical replicates; one representative experiment of *n* = 3 independent experiments shown). **(c)** Volcano plot of *Mtb* proteins differentially enriched from pull-down experiments using biotinylated ClpC1_NTD_ immobilised on streptavidin beads, incubated with AU-28, RX-76, or Cym-1 control. Fold change is calculated relative to Cym-1 for each BacPROTAC. PptT is selectively enriched by AU-28 but not RX-76. Dotted lines indicate significance thresholds (– log_10_ (*P* value) = 1.3, log₂(fold change) = ± 1). One representative of *n* = 2 independent experiments performed in triplicates shown. **(d)** Proposed model for BacPROTAC-mediated PptT engagement in cells. In a cellular environment, RX-76M cannot compete against endogenous CoA and pAp concentrations, preventing PptT engagement and degradation. In contrast, AU-28 mediates effective PptT engagement and degradation by partially displacing CoA or preferentially binding the pAp-bound form.

To further investigate BacPROTAC-mediated target engagement, we immobilised biotinylated ClpC1_NTD_ on streptavidin beads and assessed PptT pull-down from *Mtb* lysates. We found that only AU-28, and not RX-76, was able to enrich PptT (Fig. 4c), suggesting that endogenous CoA and pAp concentrations present in the lysate are sufficient to interfere with ternary complex formation by RX-76.

Collectively, these results indicate that the distinct binding modes of AU- and RX-BacPROTACs determine their susceptibility to intracellular CoA and pAp competition (Fig. 4d), providing a mechanistic basis for the differential target engagement and degradation profiles in *Mtb*. Though not an on-target PptT degrader, RX-76M likely acts through a combination of ClpC1 dysregulation and off-target Raltitrexed effects. In contrast, AU-28 represents the first validated heterobifunctional degrader of an essential mycobacterial protein, with superior antimycobacterial activity compared to the corresponding inhibitor. The integrated characterisation pipeline presented here distinguishes *bona fide* degraders from non-degraders, establishing a framework for BacPROTAC characterisation in mycobacteria.

## Discussion

In this work, we report the first heterobifunctional degrader of a native *M. tuberculosis* protein, PptT, designed by repurposing inhibitors into BacPROTACs. While previous studies have provided preliminary proof-of-concept ^4, 42, 15, 16^ none has established a true paradigm for inhibitor-to-degrader conversion. Strikingly, inhibitors with micromolar MICs against *M. tuberculosis* yielded BacPROTACs with MICs as low as 80 nM, underscoring the potency gains achievable through degrader conversion.

Alongside the BacPROTACs, we developed an integrated pipeline for their characterisation in mycobacteria, addressing the lack of a robust degrader validation workflow beyond eukaryotic systems. The limited number of validated antibodies against mycobacterial proteins precludes approaches standard for eukaryotic PROTACs,^2, 6–8^ necessitating assessment of protein degradation by quantitative proteomics. In mycobacteria, proteomic experiments are further complicated by growth stage-dependent changes in cell wall permeability and metabolic states.^43, 44^ Through systematic optimisation, we found that culturing bacteria across subinhibitory and inhibitory BacPROTAC concentrations and characterising the resulting growth inhibition phenotype by quantitative proteomics is the most effective strategy to distinguish true degraders from non-degraders. Given the resource-intensive nature of proteomics, we complemented this with MIC screening and *in vitro* degradation assays amenable to higher throughput, yielding an integrated characterisation platform.

In this study, we assessed the antimycobacterial activity of 23 BacPROTACs targeting *Mtb* PptT and prioritised compounds with improved activity relative to the parent inhibitors. However, improved whole-cell activity did not always correlate with target degradation, as illustrated by our mechanistic investigation of AU-28 and RX-76M. By combining *in vitro* and cellular assays, we validated AU-28 as the first heterobifunctional BacPROTAC degrader of a native mycobacterial protein.

This characterisation pipeline, combined with the first validated example of inhibitor-to-degrader conversion, opens new avenues for BacPROTAC-based antibiotic discovery. The thousands of *Mtb* protein ligands catalogued in databases such as ChEMBL^45^ represent a vast source of potential degrader building blocks. Converting known ligands or inhibitors into degraders could overcome limitations associated with their permeability, resistance or host-pathogen selectivity, offering unique opportunities to expand the druggable bacterial proteome. As *Mtb* treatment requires months of combination therapy with only a few drugs approved for clinical use,^20, 22^ expanding the number of druggable targets by repurposing suboptimal inhibitors into degraders represents a compelling strategy for broadening therapeutic options. This study provides the foundation for such exploration, with the aim of driving innovation against a disease that remains a leading cause of infectious disease mortality worldwide.^20^

## Methods

### Chemical synthesis

The synthesis of all compounds, including details of chemical suppliers, synthetic protocols, and characterisation data, is provided in the Supplementary Information.

### Generation of *E. coli* expression constructs

C-terminally tagged His_6_-ClpC1_NTD_, cloned in pET21a, and N-terminally tagged His_6_-PptT, cloned in pET29b, were provided by the Clausen lab. A C-terminal Avi-Tag^46^ was cloned into the pET21a-ClpC1_NTD_ plasmid, using a Q5 site directed mutagenesis kit (New England Biolabs), to obtain a ClpC1_NTD_-His_6_-AviTag (ClpC1_NTD_-Avi) construct.

Synthetic DNA codon-optimised for *E. coli* and encoding *Mtb clpP1* (Rv2461c), *clpP2* (Rv2460c) and *clpC1* (Rv3596c) was ordered from GeneArt (ThermoFisher). *clpP1* and *clpP2* genes were cloned into pET28a with C-terminal His_6_ fusions via HiFi DNA assembly (NEB), while *clpC1* was cloned into a pET49b plasmid with an N-terminal His_6_-SUMO (Smt3) fusion. Plasmid sequences were verified by nanopore sequencing.

### Protein Expression and Purification

All plasmids were transformed into *E. coli* BL21 (DE3) pLysS competent cells (Novagen #70236). Transformed cells were grown at 37°C in LB media supplemented with 50 µg/ml kanamycin (ClpP1, ClpP2, PptT, full-length ClpC1) or carbenicillin (ClpC1_NTD_, ClpC1_NTD_-Avi). Protein expression was induced upon reaching OD_600_ of 0.8 with 0.5 mM isopropyl 1-thio-βd-galactopyranoside (IPTG). Cultures were grown overnight at 18°C with agitation before cells were harvested by centrifugation. Cell pellets were flash frozen in liquid nitrogen and stored at -70°C until required for protein purification.

ClpP1 and ClpP2 cell pellets were resuspended in 50 mM HEPES pH 7.5, 300 mM NaCl, 10 mM imidazole, 10% (v/v) glycerol, 0.25 mM tris(carboxyethyl)phosphine (TCEP), supplemented with 100 U/ml benzonase nuclease (Sigma #70746-3). Cells were lysed by sonication and the crude lysate clarified by centrifugation (48,000 *g,* 45 min). The supernatant was applied to HisPur™ Ni-NTA resin (ThermoFisher #88223) and, after washing, the bound protein was eluted with 50 mM HEPES pH 7.5, 300 mM NaCl, 250 mM imidazole, 10% (v/v) glycerol, 0.25 mM TCEP. Affinity chromatography eluate was diluted in 50 mM HEPES pH 8.0, 10% (v/v) glycerol, 0.25 mM TCEP to a final concentration of 100 mM NaCl and applied to a pre-equilibrated 5 ml HiTrap Q HP anion-exchange column (Cytiva #17115401). The protein was eluted using a stepwise gradient to 1 M NaCl. Pooled fractions were concentrated and further purified by size-exclusion chromatography (SEC) on a HiLoad 16/600 Superdex 200 pg column (Cytiva #28989335) equilibrated in 50 mM HEPES pH 7.5, 300 mM NaCl, 10% (v/v) glycerol, 0.25 mM TCEP. Selected fractions were concentrated, flash frozen in liquid nitrogen and stored at -70°C for later use. Processing of the full-length ClpP1 and ClpP2 to the active ClpP1P2 complex was performed as previously described using Z-leu-leu-CHO activator dipeptide (Enzo Life Sciences #BML-PI116).^47^

Cell pellets for full-length ClpC1 purification were resuspended in 50 mM Tris-HCl pH 7.5, 500 mM NaCl, 1 mM TCEP, 10% (v/v) glycerol and lysed by sonication. Clarified lysates were purified by Ni-NTA affinity chromatography as described above. Cleavage of the N-terminal His_6_-SUMO tag was achieved by addition of Ulp1 protease to the Ni-NTA affinity eluate and overnight dialysis into 50 mM Tris-HCl pH 7.5, 300 mM NaCl, 10% (v/v) glycerol, 1 mM TCEP. Cleavage was confirmed by SDS-PAGE, and the untagged protein was collected by reverse Ni-NTA affinity chromatography. The flow-through was diluted in 50 mM Tris-HCl pH 7.5, 1 mM TCEP, 10% (v/v) glycerol to a final salt concentration of 100 mM NaCl, applied to a HiTrap Q HP column and eluted by a stepwise gradient to 1 M NaCl. SEC and flash freezing were performed as described above for ClpP1 and ClpP2 in 50 mM Tris-HCl pH 7.5, 300 mM NaCl, 1 mM TCEP, 10% (v/v) glycerol.

Cell pellets for PptT purification were resuspended in 50 mM HEPES pH 7.2, 300 mM NaCl, 10 mM MgCl₂, 10% (v/v) glycerol, 2 mM TCEP, 10 mM imidazole, 50 µM coenzyme A (CoA) (Cambridge Bioscience #CAY16147), supplemented with benzonase and lysed by sonication. Clarified lysates were purified by Ni-NTA affinity chromatography and subsequently by SEC using a HiLoad 16/600 Superdex 75 pg column (Cytiva #28989333), equilibrated with 50 mM HEPES pH 7.2, 100 mM KCl, 10 mM MgCl₂, 10% (v/v) glycerol, 2 mM TCEP.

ClpC1_NTD_ and ClpC1_NTD_-Avi were purified as previously described for ClpC1_NTD_.^4^ ClpC1_NTD_-Avi was site-specifically biotinylated using an *in vitro* biotinylation kit (Cambridge Bioscience #BPS-82511).

All protein purification steps were carried out at 4°C. Protein purity was monitored by SDS-PAGE and Coomassie staining, and correct molecular mass of purified proteins was verified by mass spectrometry.

### PptT Atto 488 Labelling

PptT was fluorescently labelled using an NHS-ester Atto 488 labelling kit (Jena Bioscience #FP-201-488). Purified PptT (90 µM) was incubated with 1 mM CoA, 100 mM sodium bicarbonate and 1 mM Atto 488 in 50 mM HEPES pH 8.0, 100 mM KCl, 10 mM MgCl_2_, 1 mM TCEP, 10% Glycerol buffer for 1 h at 15°C with shaking. Excess CoA and Atto 488 were removed by Pierce™ Dye Removal Columns (ThermoFisher #22858) and SEC (Superdex™ 75 Increase column 10/300 GL, Cytiva #29148721). Elution of labelled PptT (PptT^ATTO488^) was monitored by absorbance at 280 and 501 nm and successful labelling confirmed by SDS-PAGE and in-gel fluorescence imaging on the ChemiDoc MP imaging system (BioRad).

### Fluorescence polarisation

Fluorescence polarisation assays were performed in quadruplicate, 10 µl reaction volumes in black 384-well microplates (Corning #3821). BacPROTAC stocks were serially diluted in DMSO to give an 11-point, 2-fold dilution series resulting in final assay concentrations of 20 µM – 0.02 µM. A ClpC1_NTD_ + PptT^ATTO488^ master mix in assay buffer (50 mM HEPES pH 7.2, 100 mM KCl, 10 mM MgCl_2_, 1 mM TCEP, 10% (v/v) glycerol, 0.01% (v/v) Tween 20) was combined 9:1 with BacPROTAC dilutions to give final concentrations of 1 µM ClpC1_NTD_ and 0.1 µM PptT^ATTO488^, and a DMSO percentage of 1% (v/v). Plates were sealed, spun down (300 *g*, 1 min) and incubated at room temperature for 10 min. Seals were removed before fluorescence polarisation was measured in a CLARIOstar Plus microplate reader (BMG Labtech) at 25°C. Gains for parallel and perpendicular channels were adjusted to a target mP defined by the baseline polarisation of 1 µM ClpC1_NTD_ and 0.1 µM PptT^ATTO488^, in the absence of BacPROTAC, measured previously using a precision cuvette (Hellma #101.016-QS) and FP-8500 spectrofluorometer (Jasco). Fluorescence emission in the parallel and perpendicular channels was recorded using a fluorescein filter set (482/16 nm band-path excitation filter, 504 nm long-pass dichroic mirror and 530/40 nm band-path emission filter). For each replicate compound concentration, the mean DMSO baseline was subtracted to give the BacPROTAC-dependent Δpolarisation in mP. The mean and standard deviation of the four replicates were then plotted in GraphPad Prism version 11.0.0 (GraphPad Software).

For CoA and adenosine 3’,5’-diphosphate (pAp) competition assays, competitor compounds were serially diluted in 9% DMSO (v/v) PptT assay buffer to give a 7-point, 4-fold dilution series resulting in final assay concentrations of 320 µM – 0.078 µM. The ClpC1_NTD_, PptT^ATTO488^ master mix was split into two aliquots, one supplemented with AU-28 and the other with RX-76, each to a final assay concentration of 1.25 µM. Competitor compounds (CoA and pAp) and BacPROTAC-supplemented master mix were dispensed into the plate as described previously, to give a final well DMSO percentage of 1% (v/v). Plates were processed as described above and fluorescence polarisation values were measured with the same optics and gain settings. Polarisation data were plotted as raw polarisation values in mP and compared to BacPROTAC-only baseline (1.25 µM AU-28 / RX-76).

### *In vitro* degradation assay

*In vitro* degradation assays were performed in quadruplicate, 10 µl reaction volumes in black 384-well microplates (Corning #3821). A master mix of 1 µM PptT^ATTO488^, 0.15 µM ClpC1, 0.15 µM activated ClpP1P2 and an ATP regeneration system of 3 µM pyruvate kinase and 5 mM phosphoenolpyruvate was prepared on ice in PptT assay buffer. BacPROTAC stocks were serially diluted in DMSO to give an 11-point, 2-fold dilution series resulting in final assay concentrations of 20 µM – 0.02 µM. Compound (1 µl) and master mix (8 µl) were dispensed into assay microplates. Plates were incubated at room temperature for 10 minutes before the reaction was initiated by addition of 1 µl ATP solution (ThermoFisher #R0441) to a final concentration of 5 mM and a final well DMSO percentage of 1% (v/v). The plate was sealed with optically clear adhesive seals (ThermoFisher #AB-0580), placed in the CLARIOstar Plus microplate reader and shaken (400 rpm, 30 s) to ensure thorough mixing. The fluorescence intensity was continuously monitored (80 x 3 min cycles, 40 flashes per well, room temperature) using monochromator settings of 488/14 nm and 535/30 nm for excitation and emission, respectively. Immediately after the final read, the plate was removed and reaction mixtures of the technical replicates pooled for SDS PAGE analysis. Gels were imaged using a ChemiDoc MP Imager and the Alexa Fluor 488 in-gel fluorescence settings and subsequently stained with Coomassie Blue.

Raw fluorescence intensity data recorded on the CLARIOstar Plus plate were normalised by subtraction of t = 0 fluorescence values for each well. The mean DMSO baseline was then subtracted from each timepoint to correct for the DMSO-dependent reduction of PptT fluorescence over the assay time course. The mean and standard deviation of the four replicates were plotted in GraphPad Prism and analysed as described below.

For competition experiments with Raltitrexed, AU-08, Cym-1 and cofactors (CoA, pAp), competitor compounds were diluted to intermediate concentrations at a DMSO percentage of 9% (v/v). Master mix was prepared as above and split into two aliquots, one supplemented with AU-28 and the other with RX-76, each to a final assay concentration of 1.25 µM, and DMSO of 1% (v/v). Plates were handled and read as above with the same optics and gain settings, however data were normalised by subtraction of t = 0 only.

The final normalised Δfluorescence timepoint for each BacPROTAC concentration was plotted as a dose-response curve to compare relative BacPROTAC efficacy and potency. These concentration versus Δfluorescence plots were analysed by nonlinear regression curve fitting to determine DC50 values (concentration of half-maximal degradation) using a modified Hill equation in GraphPad Prism ([Inhibitor] vs response – variable slope (4 parameters)). For each BacPROTAC, the bottom was constrained at zero and the top at the maximum measured Δfluorescence for that BacPROTAC. Initial degradation rates (V_0_) were determined by performing a linear regression analysis over the first 15 min of the normalised Δfluorescence plots. Calculated gradients for each BacPROTAC concentration were also plotted as a dose-response. For cofactor and free ligand competition assays, initial rates were calculated from the t = 0 normalised plots.

### Mycobacterial culture

Liquid cultures of *M. tuberculosis* (*Mtb*) H37Rv were grown in Middlebrook 7H9 medium (Merck #M0178) supplemented with 10% (v/v) ADC (BD #211887), 0.1% (v/v) Tween80 (Sigma Aldrich #P4780) and 0.4% (v/v) glycerol, at 37°C with agitation. Solid cultures of *Mtb* H37Rv were grown on Middlebrook 7H10 medium (Merck #M0303) supplemented with 10% (v/v) OADC (BD #211886), 0.1% (v/v) Tween80 and 0.5% (v/v) glycerol, at 37°C. As required, growth media were supplemented with kanamycin up to 25 µg/ml and anhydrotetracycline (ATc) up to 100 ng/ml.

### MIC determination

The minimum inhibitory concentration (MIC, IC_90_) was determined via broth microdilution. Briefly, DMSO stocks of each compound were diluted in a 10-point, 2-fold dilution series at 100-fold the final assay concentration, yielding final assay concentrations of 20–0.04 µM. *Mtb* cultures were grown at 37°C with agitation until reaching exponential phase, up to OD_600_ = 1, before dilution. In a 96-well plate (Falcon #353219), 1 µl of either compound, DMSO (growth control) or 3 mM rifampicin (no-growth control) were diluted in a final volume of 100 µl with cell suspension at OD_600_ of 0.005. Four technical replicates were carried out per compound. Plates were incubated at 37°C for 10 days before OD_600_ was measured using the BioTek Synergy H1 Multimode Reader (Agilent). Relative viability was calculated relative to the growth (DMSO, 100%) and no growth (rifampicin, 0%) controls. MIC is defined as lowest concentration with at least 90% growth inhibition.

### Global proteomics sample preparation

Compound dilutions were prepared in DMSO, and 5 µl of either compound or vehicle control (DMSO) was dispensed in triplicate into 48-well plates (Falcon #353230). Across repeats, *Mtb* was treated with AU-28 at concentrations between 1.25-0.01 µM, AU-08 and Cym1 at 2.5-0.04 µM and RX-76M at 2.5-0.04 µM. *Mtb* cultures were grown at 37°C with agitation until reaching an OD_600_ of 0.2-0.3. To each well, 495 µl of this cell suspension was dispensed and plates were incubated static at 37°C for either 6, 24 or 72 h. Following incubation, the treated cell suspension was centrifuged (4,000 *g* 3 min). The cell pellet was washed in PBS. After harvesting, the cell pellet was resuspended in 100 µl lysis buffer (50 mM Tris-HCl pH 8, 150 mM NaCl, 0.5% (w/v) sodium deoxycholate, 2% (w/v) SDS, containing cOmplete™ EDTA-free Protease Inhibitor Cocktail (Roche #11836170001)). *Mtb* was heat inactivated by incubating at 95°C for 20 min. Cells were transferred to 1.5 ml Bioruptor Pico Microtubes (Diagenode #C30010016) and lysed using the Bioruptor Pico (Diagenode: 4°C, 10 cycles of 30 s on, 30 s off), followed by centrifugation (4,000 *g*, 3 min) to clarify the lysates. Protein concentration was determined via BCA assay (ThermoFisher #A55861), and concentrations were normalised in lysis buffer.

Clear lysates were adjusted to 2-5% SDS (w/v) then purified using PROTIFI™ S-Trap™ 96-well plates or micro spin columns (ProtiFi) according to manufacturer’s protocol. Briefly, SDS-containing samples were reduced, alkylated then acidified with 20 mM DL-Dithiothreitol (Merck #D0632), 40 mM Iodoacetamide (Merck #I1149) and 1.2% Phosphoric acid (v/v) (Merck #49685) respectively (final concentrations). The acidified lysates were diluted with 5-6x Binding Buffer (100 mM TEAB pH 8.5, 90% MS-grade MeOH) and applied to the S-Trap™ plate columns. Bound proteins were washed 3x with Binding Buffer, then digested with MS-grade Trypsin (ThermoFisher #90057) in Digestion Buffer (50 mM TEAB, pH 8.5) at 37°C overnight or 47°C for 2 h, at S:E ratio of 25 (μg/μg), with a minimum of 1 μg/well. Digested peptides were eluted sequentially with Digestion Buffer, Elution Buffer 1 (0.1% formic acid (FA)) and Elution Buffer 2 (50% acetonitrile (ACN), 0.1% FA) to reach 10-11% ACN in the final elution. The elution was dried into powder with SpeedVac, before LC-MS dried peptides were reconstituted in 50 μl Elution Buffer 1.

### Preparation of *Mtb* lysates and pull-down experiments

*Mtb* cultures grown to exponential phase were centrifuged (3,200 *g*, 10 min). The cell pellet was washed in PBS-0.05% (v/v) Tween80, followed by centrifugation. The cell pellet was resuspended sequentially in 1/2 and then 1/50^th^ the original culture volume in PBS. The concentrated cell suspension was centrifuged (21,000 *g*, 5 min) and resuspended in lysis buffer (50 mM HEPES, 150 mM KCl, 10 mM MgCl_2_, 5% (v/v) glycerol, 0.25 mM TCEP, containing cOmplete™ EDTA-free Protease Inhibitor Cocktail). Cells were transferred into Lysing Matrix B tubes (MP Biomedicals™ #116911100) and lysed by mechanical disruption using the FastPrep®-24 bead beater (MP Biomedicals™: 3 cycles of 60 s at 6.5 m/s) with incubation on ice between cycles. The lysate was clarified by centrifugation (21,000 *g*, 20 min). The clarified lysate was twice passed through 0.22 µm filter spin columns (Corning #8160), and then flash-frozen in liquid nitrogen and stored at -70°C.

MagReSyn® Streptavidin MS (ReSyn #MR-STP002) beads were equilibrated in lysis buffer. Biotinylated ClpC1_NTD_-Avi was incubated with the MagReSyn beads. Compounds were diluted to 10 µM at 1% (v/v) DMSO in the lysis buffer, and incubated with the protein-conjugated MagReSyn beads. Samples were split into technical triplicates before adding lysate equivalent to 250 µg total protein. Following lysate incubation, samples were washed with 50 mM TEAB pH 8.5.

Samples were reduced and alkylated with 5 mM TCEP (Santa Cruz Biotechnology #sc-203290A), 10 mM Chloroacetamide (ThermoFisher #16336562) in 50 mM TEAB pH 8.5. Proteins were digested overnight at 37°C with 0.1 μg/μl Trypsin/LysC (ThermoFisher #A41007) in 50 mM TEAB pH 8.5. Samples were acidified by addition of 10% FA to 1% final concentration.

### Mass spectrometry analysis by LC-MS

Approximately 200 ng of samples from both global and pull-down proteomics were loaded onto Evotips, prepared according to the manufacturer’s protocol. The samples were analysed on an Evosep One LC system (Evosep) coupled with a timsTOF Pro2 mass spectrometer (Bruker) via a CaptiveSpray nano-electrospray ion source. Data for all samples was acquired in diaPASEF mode using the 60 SPD predefined method on Evosep One, which was fitted with an 8 cm column (Evosep #EV1109). Mobile phase A was 0.1% formic acid in water and mobile phase B 0.1% formic acid in acetonitrile. Mass spectra were acquired from 100 to 1700 m/z. The ion mobility range was set to 0.6–1.60 Vs cm-2. TIMS accumulation and ramp times were set to 100 ms. 12 diaPASEF scans were collected per one TIMS-MS scan, giving a duty cycle of 1.37 s. 24 mass windows were set over the mass range 262.2–1199.6m/z and mobility range 0.6–1.60 Vs cm-2. The collision energy was increased linearly from 20 eV to 59 eV between 0.6 and 1.60 Vs cm-2.

Raw data was searched using Biognosys Spectronaut® against MTb H37Rv (Taxon ID: 83332) UniProt proteome (UP000001584) and contaminants FASTA files. BGS factory settings were used with QUANT2.0 for protein LFQ Method, global median normalisation strategy with automatic row selection was performed. Missing values were imputed using run wise imputing. Quantification was performed at the MS2 level.

### Proteomic data and statistical analysis

Protein quantity data was analysed and visualised with custom python scripts (https://github.com/yjzhng/omicViz). Compound-treated samples were compared against vehicle controls by unpaired two-tailed t-tests. P-value was used without adjustment due to sample size and effect size limit. Significance of differential expression was determined with default thresholds (absolute log2 fold change > 1 and -log10 P-value > 1.3) unless stated otherwise. For comparison between compounds, t-tests were performed and significance was determined as above but with an additional criterion that the protein must also be significant in either compound alone (vehicle-normalised).

### Generation of CRISPRi knockdown strains

Candidate sgRNA was identified downstream of a protospacer adjacent motif on the non-template strand of *pptT*, and RxnReady oligos (IDT) were ordered (5’-GGGAGCTGATCGCATCCAGCACACCAT-3’, 5’- AAACATGGTGTGCTGGATGCGATCAGC-3’). Oligos were phosphorylated, annealed and ligated into the BsmBI-v2-digested CRISPRi plasmid pLJR965. Plasmid sequence was verified by nanopore sequencing.

The *pptT* knockdown plasmid (pLJR965-PptT) and the negative control plasmid containing a non-targeting sgRNA (pLJR965) were transformed into *Mtb*.^48^ Electrocompetent *Mtb* cells were generated from culture grown to OD_600_ ∼1. The culture was harvested by centrifugation (3,000 *g*, 5 min), and washed 3 times in 10% (v/v) glycerol in sequentially reducing volumes before final resuspension in 1/20^th^ the original culture volume. For electroporation, 100 ng of plasmid DNA was mixed with 200 µl electrocompetent cells in a 0.2 cm electroporation cuvette (BioRad #1652086), and electroporation was carried out using Gene Pulser Xcell Electroporation System (BioRad #1652660: 2,500 V, 1000 Ω and 25 μF). Bacteria were recovered overnight in 1 ml media, before plating onto 7H10 supplemented with 25 µg/ml kanamycin and incubation at 37°C.

### Checkerboard assay

Using the checkerboard method, a two-dimensional array of serial dilutions of ATc and compound were prepared along each axis of 96-well plates. A dilution series of ATc was prepared to 100-fold final assay concentrations in a 10-point, 2-fold dilution series in 70% ethanol (v/v), to yield final assay concentrations of 100 – 0.2 ng/ml. Compounds in DMSO were diluted in a 7-point, 2-fold dilution series at 100-fold the final assay concentration, yielding the indicated concentrations (0.63 – 0.01 µM for AU-28 and 5.0 – 0.08 µM for RX-76M). Cultures of *Mtb* were grown until reaching exponential phase, before dilution. In the assay plates, 1 µl of each ATc and compound (or respective vehicle control) were diluted in a final volume of 100 µl with the respective cell suspension at OD_600_ of 0.005 in 7H9, supplemented with 6.25 µg/ml kanamycin in the presence of the CRISPRi plasmid. Growth (DMSO) and no growth (rifampicin) controls were prepared as above. At least 3 technical replicates were carried out for each condition. Plates were incubated at 37°C for 10 days before measuring OD_600_ using the BioTek Synergy H1 Multimode Reader. Relative viability was calculated relative to the growth (1% (v/v) DMSO + 0.7% (v/v) ethanol, 100% growth) and no growth (30 µM rifampicin, 0% growth) controls.

### Cytotoxicity assay

Cytotoxicity of each compound was assessed using the HepG2 human hepatocellular carcinoma cell line. Compounds were dispensed into 384-well microplates (Greiner # 781976) across an 8-point, 3-fold dilution series (top concentration 50 µM) using an Echo 650 acoustic liquid handler. Wells were back-filled with DMSO to maintain a final DMSO concentration of 0.5% (v/v). Cells were then seeded at 5,000 (24 h) or 3,500 (48 h) cells per well using an Integra Viafill bulk dispenser (50 µl per well) and incubated for 24 or 48 h at 37°C with 5% CO₂. Following incubation, cells were fixed with 4% paraformaldehyde (Merck #F8775), permeabilised with 0.2% Triton X-100 (Merck #T8787), and stained with 2 µg/ml DAPI (Merck #10236276001) to visualise nuclei. Nuclei were imaged at 377 nm excitation and 447 nm emission using a Celigo image cytometer and quantified using the Direct Cell Count module in CeligoPro software (v5.5.1.0). All conditions were performed in triplicate. Staurosporine (Cambridge Bioscience #S7600) and DMSO alone served as positive and negative controls, respectively. Heat maps in Extended Data Fig.1 were generated in Python using mean viability values from three technical replicates.

## Acknowledgements

We thank all members of the Morreale lab for experimental advice and helpful discussions; Joanna Evans and Julio Ortiz Canseco for H37Rv protocols and experimental advice; the Scientific Technology Platforms (STPs) at the Francis Crick Institute, particularly Mike Howell, Ok-Ryul Song, Isabel Gameiro-Ros and Scott Warchal (High-Throughput Screening STP) and Manuela Natoli (Cell Sciences) for the cytotoxicity assays; Joanna Redmond and Christelle Soudy (Chemical Biology STP); Aini Vuorinen (Proteomics STP) for advice on proteomics experimental design; Geoff Kelly for assistance with NMR data collection at the MRC Biomedical NMR Centre; and Tim Clausen for providing plasmids. We thank Alessio Ciulli, Eachan Johnson, Maximiliano Gutierrez, Andrea Testa, and Katrin Rittinger for helpful discussions.

## Author contributions

J.D., S.F., D.B. and F.E.M. designed the experiments. J.D. performed the chemical synthesis of the BacPROTACs and control compounds. S.F. performed experiments with *M. tuberculosis* including MIC assays, CRISPRi, lysate preparation and pull-down experiments. D.B. performed *in vitro* binding and degradation assays with support from S.K.. S.F. and Y.Z. prepared samples for quantitative proteomics and analysed the data with support from L.K. and M.S.. F.E.M. conceived and coordinated the research project, and prepared the manuscript together with J.D., D.B. and S.F., with input from all authors. J.D., D.B. and S.F. contributed equally and are listed in a randomised order.

## Materials & Correspondence

Correspondence to Francesca E. Morreale

## Data availability

The mass spectrometry proteomics data have been deposited to the ProteomeXchange Consortium via the PRIDE ^49^ partner repository with the dataset identifier PXD079592. Full, uncropped images of all gels are provided in Supplementary Fig. 1. Materials are available upon reasonable request under material transfer agreements (MTA) with the Francis Crick Institute.

---

**Extended Data Figure 1.**
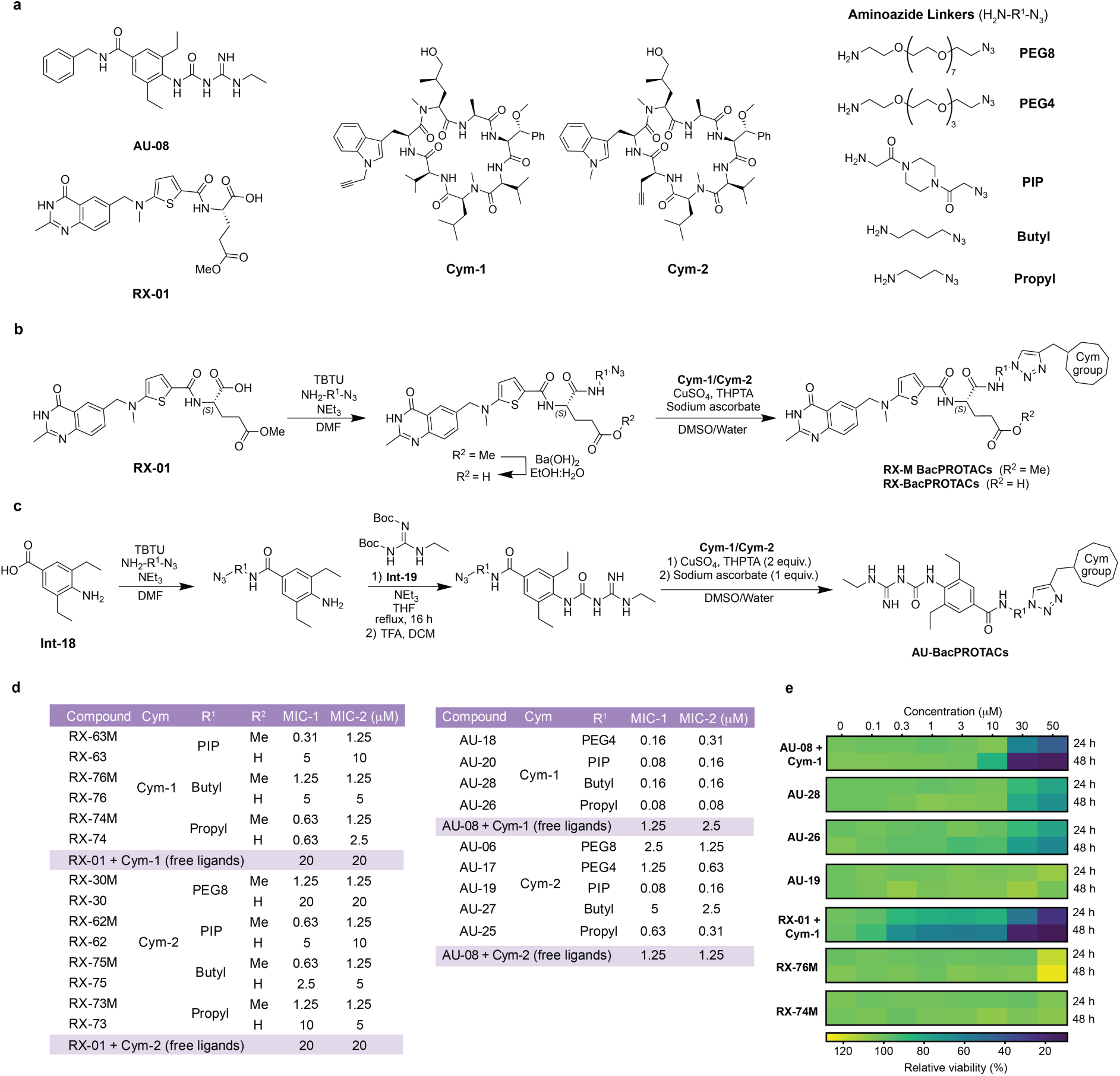
BacPROTAC synthesis and cellular activities. **(a)** Chemical structures of free ligands AU-08 and RX-01 (PptT ligands), Cym-1 and Cym-2 (ClpC1 ligands) with different exit vectors as alkyne click handles, and aminoazide linkers. **(b)** Synthetic scheme for RX-BacPROTACs. The linker is attached to RX-01 via amidation. Methyl ester variants (R^2^ = Me) are reacted with a cyclomarin derivative via click chemistry to afford RX-M BacPROTACs. Alternatively, the methyl ester can be hydrolysed by Ba(OH)₂ to give the carboxylic acid (R₂ = H), which can undergo a click reaction with the cyclomarin derivative to afford RX BacPROTACs. **(c)** Synthetic scheme for AU-BacPROTACs. The linker is attached to aminobenzoic acid Int-18 before introduction of the amidinourea moiety using Int-19, followed by Boc deprotection (TFA, DCM). A subsequent click reaction with a Cym derivative affords the AU-BacPROTACs. **(d)** Table of MIC values obtained from *n* = 2 independent experiments (MIC-1 and MIC-2) performed in quadruplicate, where the synthesised BacPROTACs are tested against *Mycobacterium tuberculosis* H37Rv. MIC-1 is plotted in Fig. 1f. MIC is defined as the lowest concentration inhibiting bacterial growth by 90% relative to DMSO (100% growth) and rifampicin (0% growth) controls. **(e)** Heatmap of HepG2 cell viability following treatment with BacPROTACs or free ligands at the indicated concentrations for 24 and 48 h (*n* = 2 independent experiments performed in technical triplicate). Cell viability is expressed as a percentage relative to DMSO controls (100% viability). Free ligands show greater cytotoxicity than the corresponding BacPROTACs, which are well tolerated at the tested concentrations.

**Extended Data Figure 2.**
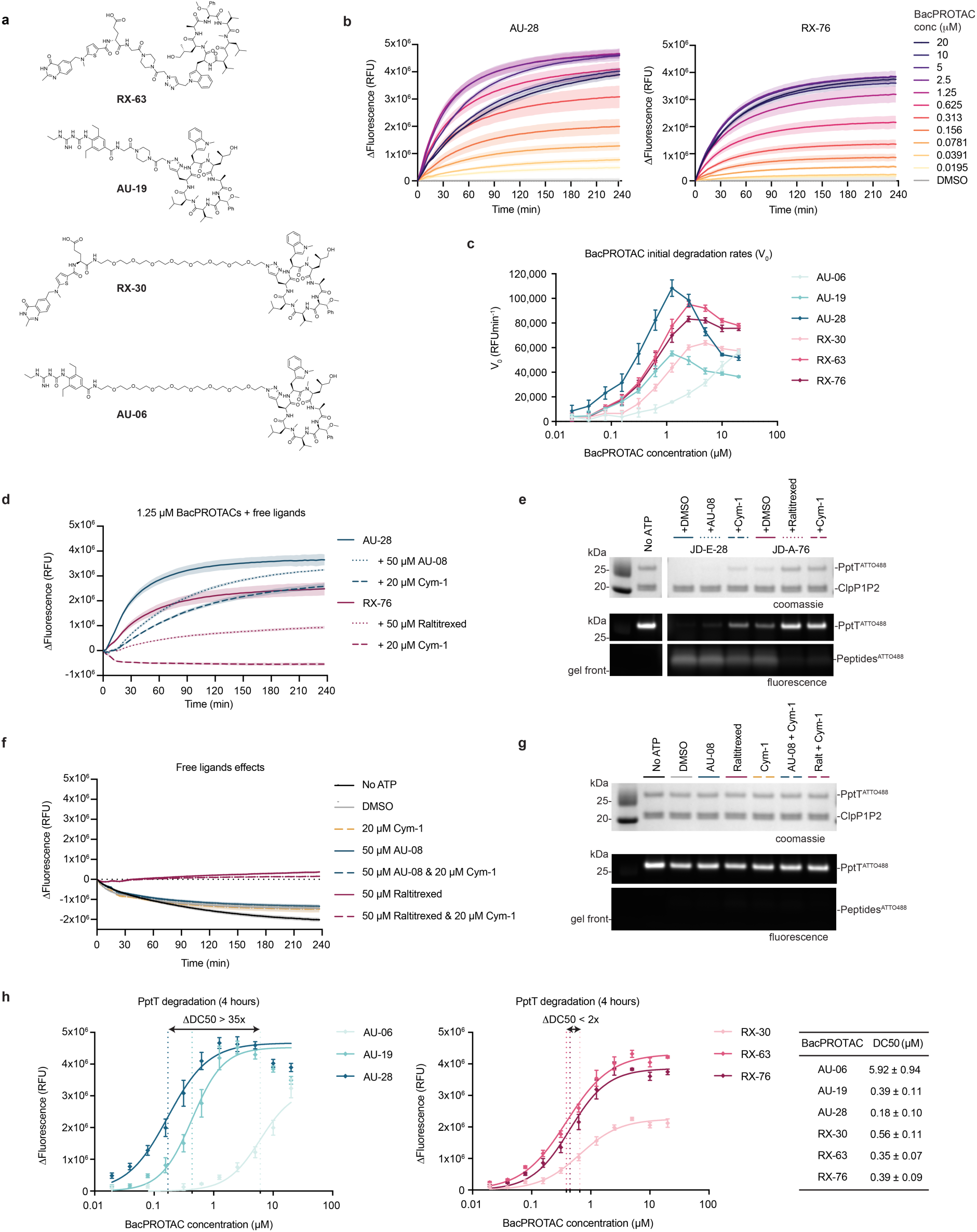
*In vitro* characterisation of BacPROTAC- induced PptT degradation. **(a)** Chemical structures of RX-63, AU-19, RX-30 and AU-06 BacPROTACs tested *in vitro* alongside AU-28 and RX-76 (shown in Fig. 2a). **(b)** Kinetic plots of PptT^ATTO488^ degradation by AU-28 and RX-76 at increasing concentrations. The corresponding fluorescence intensity endpoint (t = 4 h) is plotted in Fig. 2e. Fluorescence intensity values are normalised by subtraction of t = 0 and DMSO background. Data are mean ± s.d. (shaded area) from *n* = 4 technical replicates. **(c)** Initial PptT degradation rates across all tested BacPROTAC concentrations, estimated by linear regression on the first 15 minutes of the normalised kinetic curves and plotted as a function of concentration. **(d)** Kinetic plots of PptT^ATTO488^ degradation by 1.25 µM AU-28 and RX-76 in the presence of excess free ligand, showing a reduction in the rate and extent of degradation. Fluorescence intensity values are normalised by subtraction of t = 0 and data are mean ± s.d. (*n* = 3 technical replicates). **(e)** SDS-PAGE analysis of the fluorescence degradation assay endpoint (t = 4 h) shown in (d), imaged by in-gel fluorescence and Coomassie staining, corroborating the results observed by fluorescence intensity readout. **(f)** Fluorescence intensity readout from incubating PptT^ATTO488^ with the free ligands, showing no degradation. The downward trend, likely resulting from gradual fluorescence quenching due to partial unfolding or aggregation, is commonly observed for PptT^ATTO488^. Saturating Raltitrexed concentrations appear to stabilise the protein against this effect. These controls validate that BacPROTAC-mediated induced proximity is required for effective degradation. **(g)** SDS-PAGE analysis of the free ligand degradation assay endpoint (t = 4 h) shown in (f), imaged by in-gel fluorescence and Coomassie staining, confirming the absence of fluorescent peptides usually derived from ClpC1P1P2-mediated PptT degradation. **(h)** Curve fitting of concentration-dependent PptT degradation by BacPROTACs shown in Fig. 2e. Data are mean ± s.d. of *n* = 4 technical replicates; curves are four-parameter logistic (4PL) fits constrained at the bottom and top to zero and the measured maximum, respectively. Calculated half-maximal degradation concentrations (DC50) are shown to the right of the plots as the mean ± s.d. from *n* = 3 independent experiments. Data in (b), (c) and (h) are mean ± s.d. (*n* = 4 technical replicates; one representative experiment of *n* = 3 independent experiments shown).

**Extended Data Figure 3.**
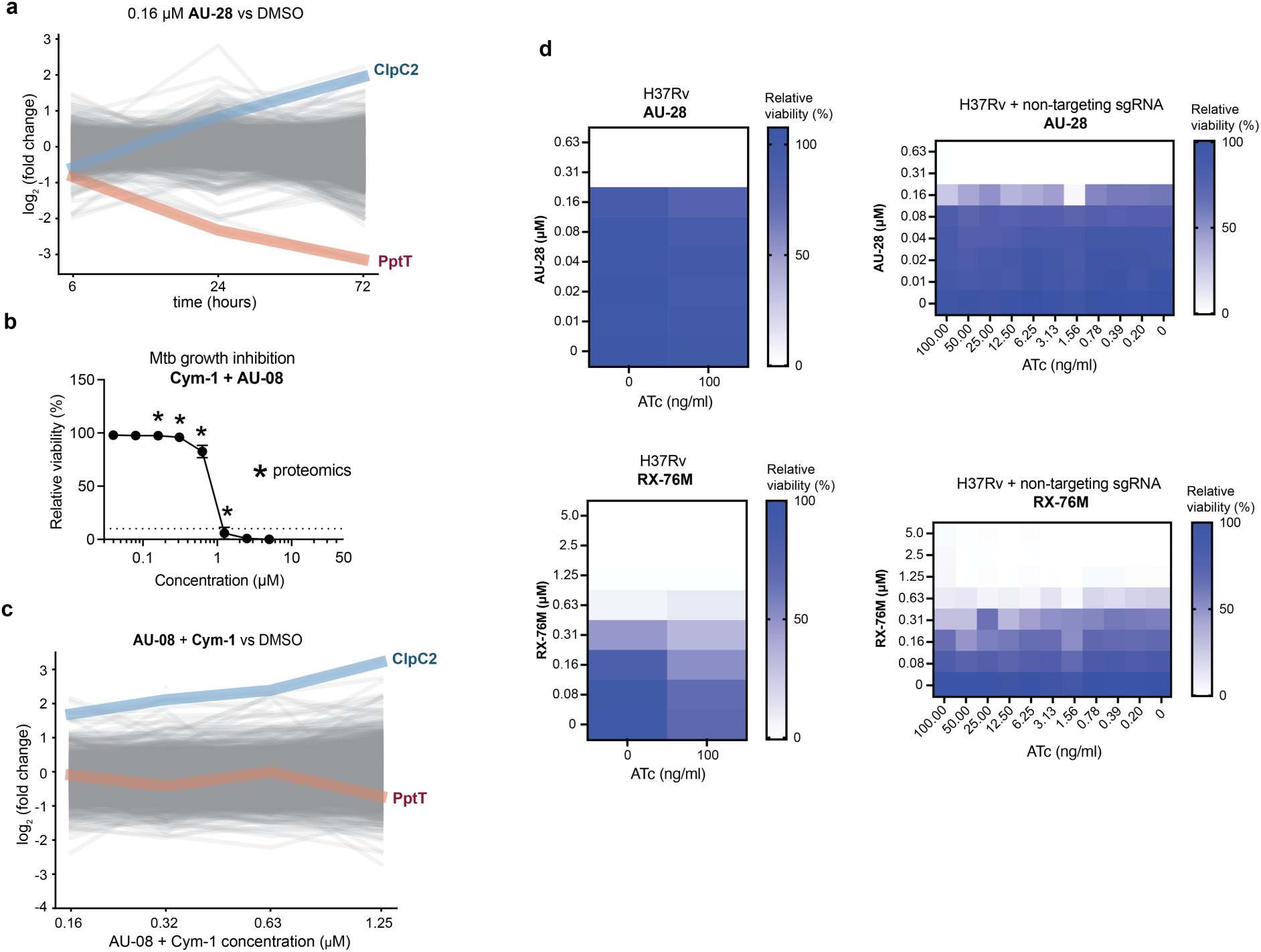
Cellular characterisation of BacPROTAC and control compounds. **(a)** Proteomic changes across three time points following treatment of Mtb cultures with 0.16 µM AU-28, showing time-dependent PptT degradation and ClpC2 upregulation (72 h: *n* = 3; 6 h and 24 h: *n* = 1 performed in technical triplicate). **(b)** Mtb growth inhibition induced by the combination of AU-08 and Cym-1 free ligands, corresponding to the AU-28 BacPROTAC. The dotted line indicates the minimum inhibitory concentration (MIC) threshold (90% growth inhibition), relative to DMSO (100% growth) and rifampicin (0% growth) controls. Asterisks indicate concentrations selected for proteomic analysis. Data are mean ± s.d. (representative of *n* = 4 independent experiments performed in technical triplicate). **(c)** Proteomic changes across concentrations of AU-08 and Cym-1 free ligand combination (*n* = 3 biological replicates, each performed in triplicate), showing concentration-dependent ClpC2 upregulation but no PptT degradation, confirming that PptT degradation observed for AU-28 (Fig. 3c,d) requires bifunctional engagement. **(d)** Control experiments associated with the CRISPRi-mediated transcriptional repression of *pptT* in *Mtb* modulated by increasing ATc concentrations shown in Fig. 3f. Heat maps show a checkerboard assay in which ATc and BacPROTAC are co-titrated in wild-type *Mtb* H37Rv and in H37Rv transformed with a non-targeting sgRNA, confirming that ATc-dependent hypersensitisation to AU-28 requires the presence of dCas9 and *pptT*-targeting guide RNA. *Mtb* growth inhibition is normalised to vehicle (100% growth) and rifampicin (0% growth) controls. Data are mean of *n* = 3 technical replicates; one representative of *n* = 2 independent experiments shown.

**Extended Data Figure 4.**
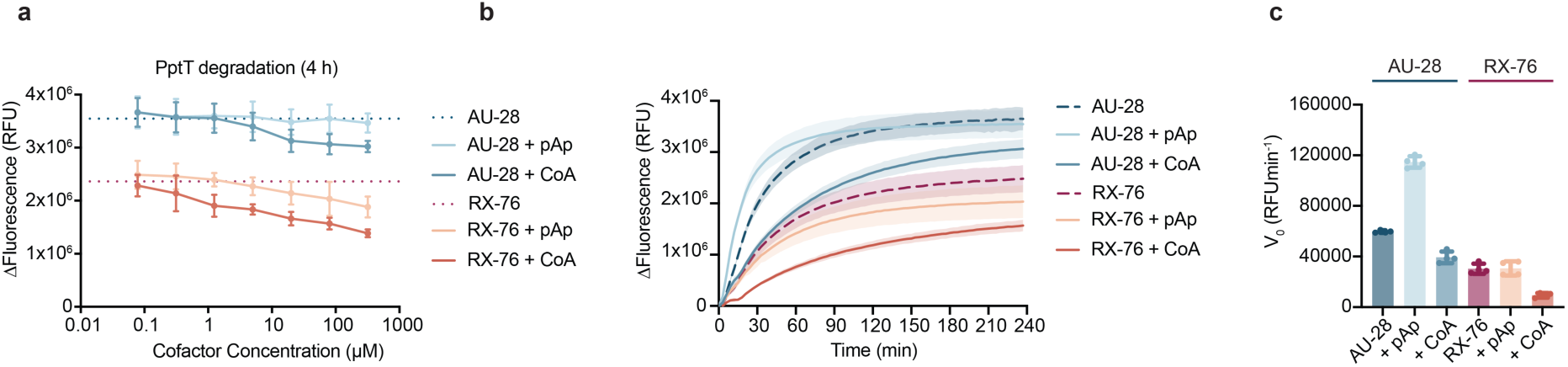
CoA and pAp modulate BacPROTAC-mediated PptT degradation. **(a)** PptT ^ATTO488^ degradation induced by 1.25 µM AU-28 or RX-76 in the presence of increasing CoA or pAp concentrations, quantified as the difference in fluorescence intensity between t = 0 and the assay endpoint (t = 4 h). Both CoA and pAp inhibit RX-76-induced PptT degradation, whereas only CoA partially inhibits AU-28, which retains higher maximal degradation. **(b)** Kinetic plots of PptT^ATTO488^ degradation by 1.25 µM AU-28 or RX-76 in the presence of 80 µM CoA or pAp. Fluorescence intensity values are normalised by subtraction of t = 0. **(c)** Initial PptT degradation rates shown as a bar plot, estimated by linear regression on the first 15 minutes of the normalised kinetic curves in (b). Data in **(a, b, c)** are mean ± s.d. (*n* = 4 technical replicates; one representative experiment of *n* = 3 independent experiments shown).

